## Supplemental Figures S1-6 and Table 1 for "Periaqueductal gray activates antipredatory neural responses in the amygdala of foraging rats"

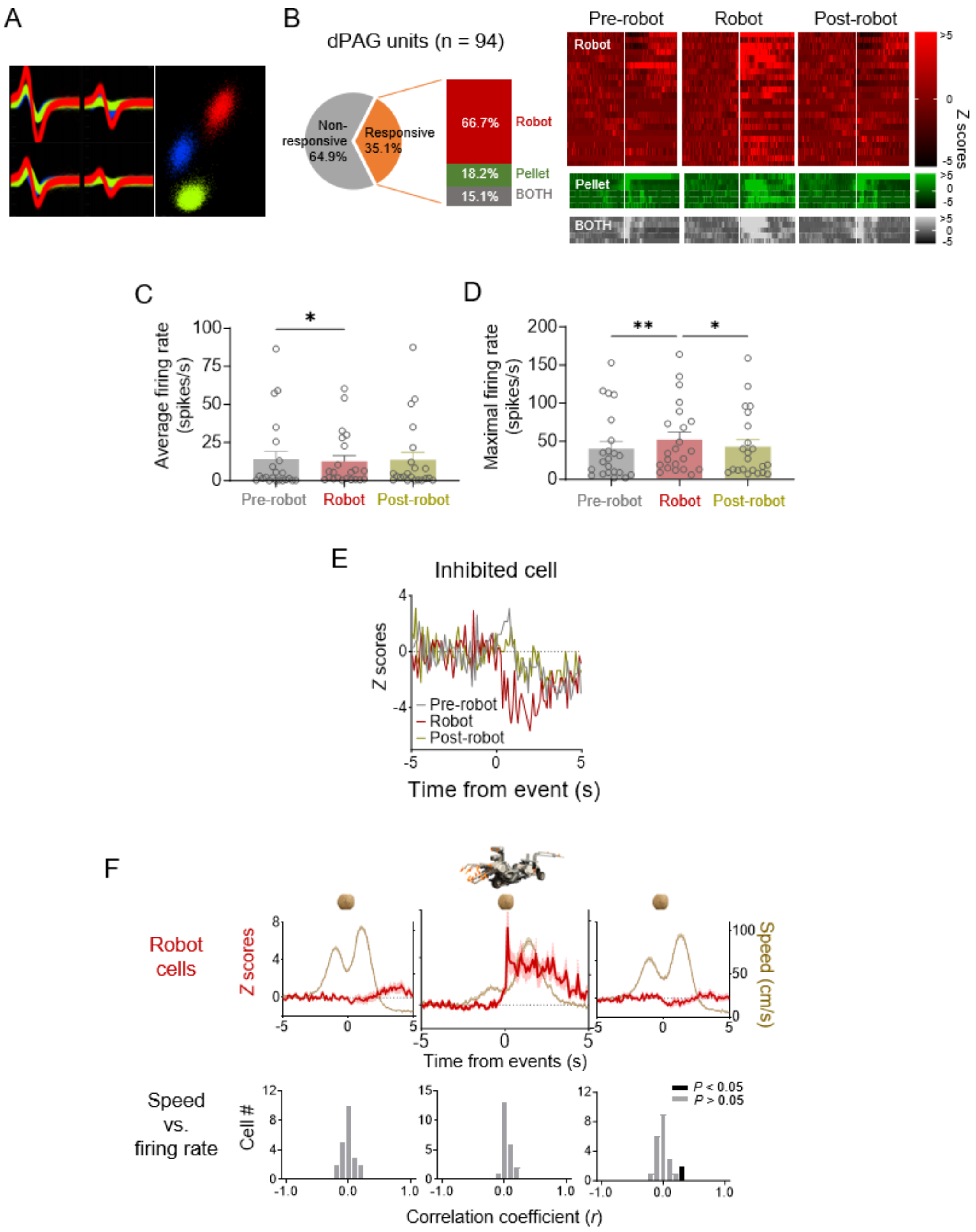


**Figure S1. dPAG unit cell types.** (A) Representative multiple single units recorded in the dPAG. (B) A subset (35.1%) of dPAG neurons showed increased firing rates in response to the robot, food pellet, or both, with 66.7% responding exclusively to the robot. Units were categorized based on z-score responses: "robot cells" for z > 3 during robot phase and z < 3 during pre-robot phase; "pellet cells" for z > 3 during pre-robot phase and z < 3 during robot phase; and "BOTH cells" for z > 3 in both phases. Raster plots display activity of robot, pellet, and BOTH cells across pre-robot, robot, and post-robot sessions. (C, D) Average (C) and maximum (D) firing rates of dPAG robot cells during pre-robot, robot, and post-robot sessions. (E) One neuron showed decreased firing rates ($z < -3$) in response to the robot during the robot phase. (F) Top row: Mean firing rate (± SEM; shaded areas) of robot cells and movement speed (± SEM; shaded areas) of animals across sessions. Bottom row: Correlation coefficients between firing rate and movement speed for the robot cell group during each session.

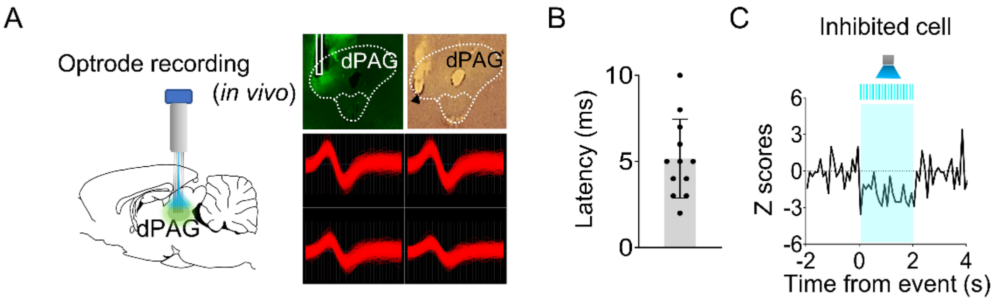


**Figure S2. Optrode recording in the dPAG.** (A) Virus injection and optrode implantation targeted at the dPAG (*left*). Photomicrographs display virus expression, optic fiber/electrode tips, and a representative dPAG neuron (*right*). Light stimulation was applied during single unit recordings in anesthetized rats. (B) Response latencies of stimulation-responsive dPAG cells. (C) A single dPAG cell exhibited inhibited responses to optical stimulation of the dPAG.

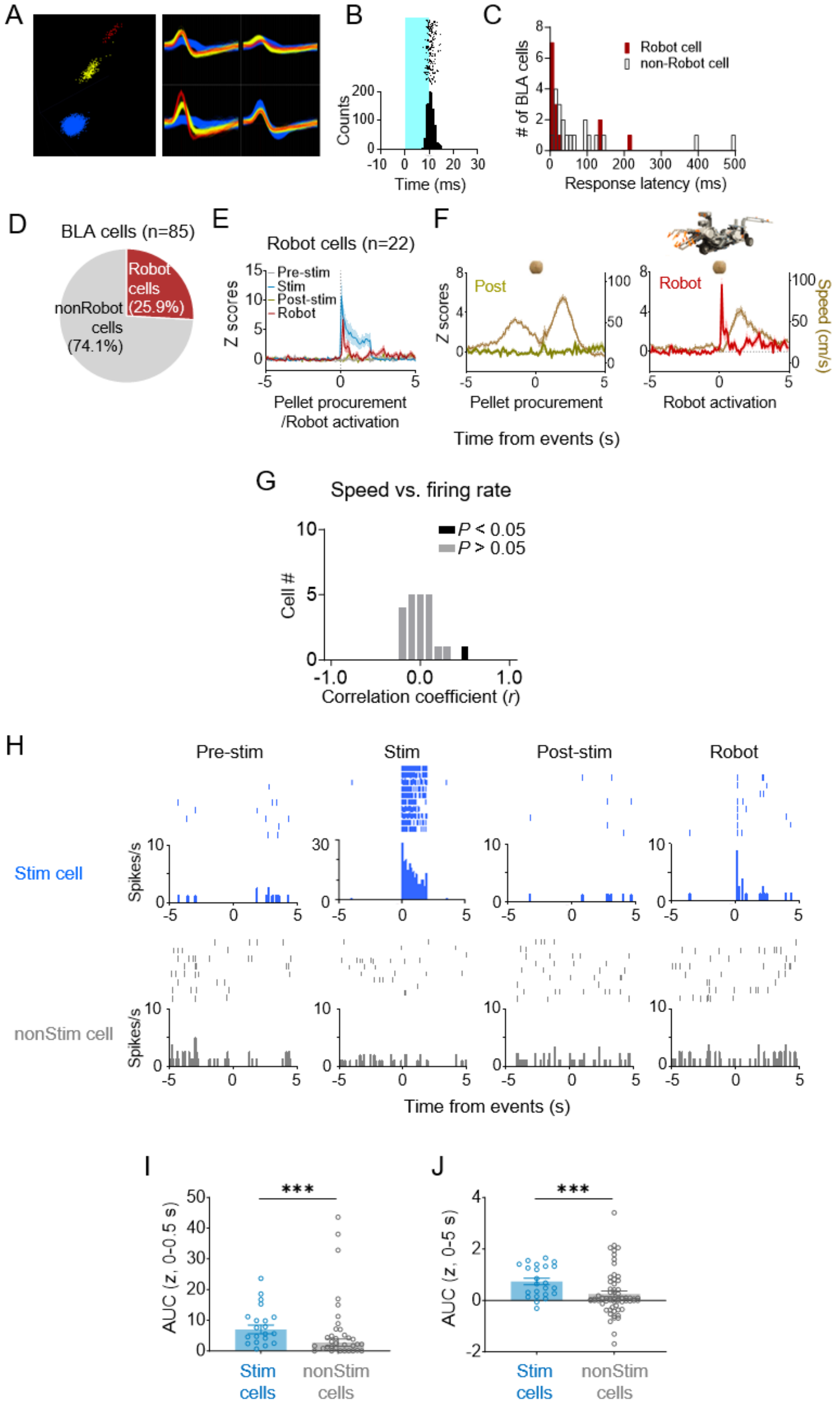


**Figure S3. Characteristics of BLA cells.** (A) Representative waveforms of BLA cells.

(B) A representative raster plot and peri-event time histograms for 20-Hz light stimulations (10-ms pulse width, 2-s duration). (C) Response latencies of BLA cells to dPAG stimulation. (D) Out of 85 units from the BLA, 25.9% immediately responded to the robot. (E) A pre-event time histogram aligning BLA activity with pellet procurement (pre-stim and post-stim sessions), dPAG stimulation (stim session), or robot activation (robot session). (F) Firing rates of robot cells and movement speeds during pellet procurement (post-robot session) and robot activation (robot sessions). (G) All but one BLA cell showed no significant correlation between firing rate and movement speed. (H) Representative raster plots and peri-event time histograms for a dPAG stimulation-responsive BLA cell reacting to the robot predator (*top*; also see Figure 3I, left column) and a dPAG stimulation-nonresponsive BLA cell (*bottom*; also see Figure 3I, right column). (I, J) dPAG stimulation-responsive units showed higher levels of robot-evoked firings than stimulation-nonresponsive units during both short (I; 0-0.5 s post-robot activation; $U$ = 339.0, $P$ = 0.0001; Mann-Whitney U test) and extended (J; 0-5 s post-robot activation; $U$ = 384.5, $P$ = 0.0009; Mann-Whitney U test) periods. *** denotes $P < 0.001$ compared to the non-stim pairs.

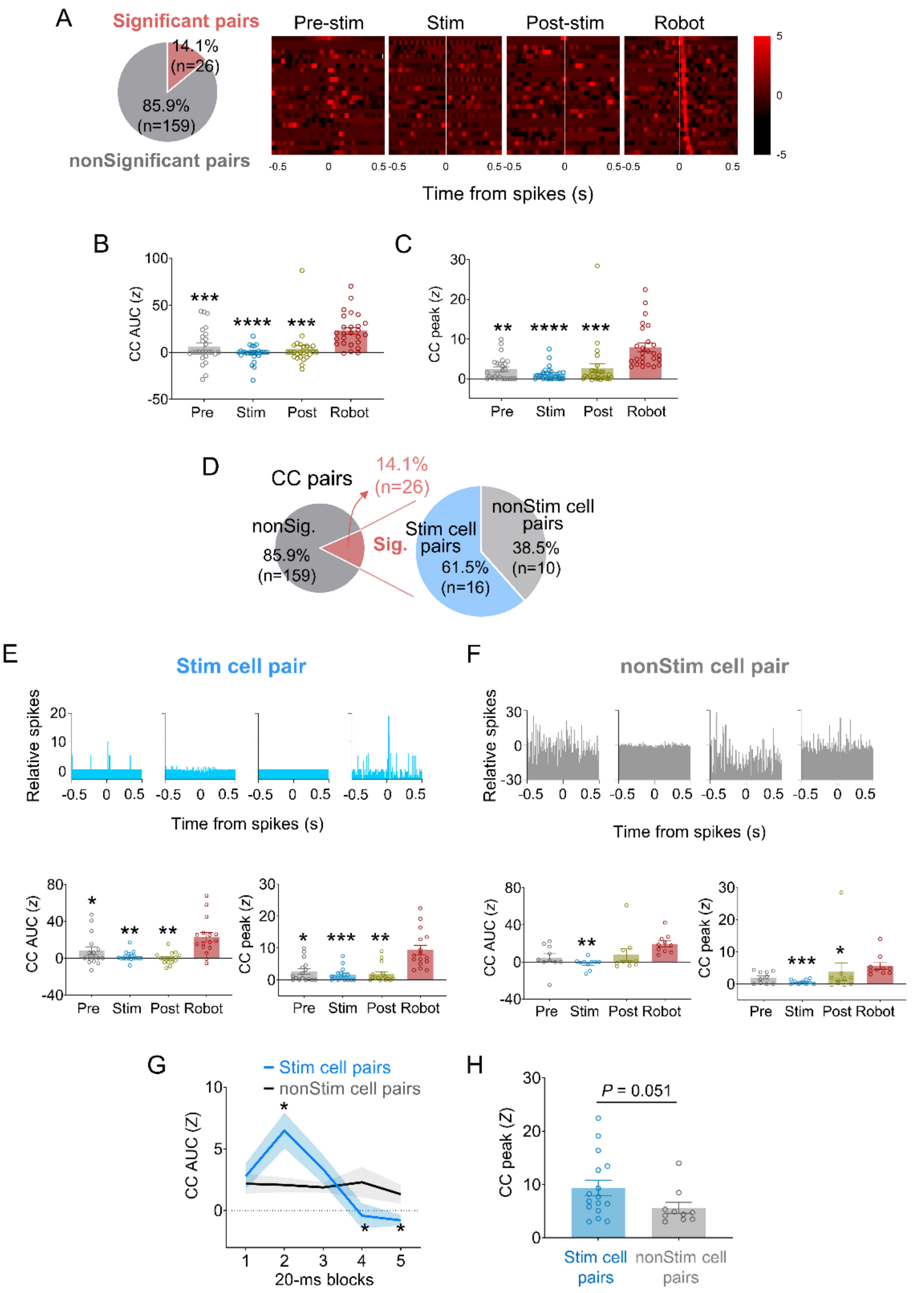


**Figure S4. Spike synchrony in dPAG-stimulated BLA neurons under predatory threats.** (A) Of 185 CCs from simultaneously recorded BLA neuron pairs, 14.1% exhibited significant peaks ($z > 3$) during robot sessions. (B, C) Mean CC AUC (B) and

CC peak values (C) increased during robot sessions compared to other sessions ($Z$s $> 29.36$, $P < 0.0001$, Friedman test; $P$s $< 0.01$ for pre-stimulation vs. robot, stimulation vs. robot, and post-stimulation vs. robot sessions comparisons, Dunn’s test). **, ***, and **** denote $P < 0.01$, $P < 0.001$, and $P < 0.0001$ compared to robot session, respectively. (D) Among BLA cell pairs with significant synchrony during robot sessions, 61.5% included dPAG-stimulated BLA cells (stim pairs) and 38.5% did not (non-stim pairs). (E) Stim pairs (*top*, a representative pair) showed enhanced spike synchrony during robot sessions, with higher mean CC AUC (*bottom left*) and peak (*bottom right*) values compared to other sessions ($Z$s $> 17.42$, $P < 0.001$, Friedman test; $P$s $< 0.037$ for pre-stimulation vs. robot, stimulation vs. robot, and post-stimulation vs. robot sessions comparisons, Dunn’s test). *, **, and *** denote $P < 0.05$, $P < 0.01$, and $P < 0.001$ compared to the robot session, respectively. (F) Non-stim pairs (*top*, a representative pair) also displayed significant synchrony during robot sessions, with no difference in CC AUC (*bottom left*) between robot and pellet-only sessions, but higher peak area during robot vs. stimulation session ($Z = 13.84$, $P < 0.01$, Friedman test; $P = 0.0017$ for the stimulation vs. robot sessions comparison, Dunn’s test). CC peak (*bottom right*) during robot session was higher than during stimulation and post-stimulation sessions ($Z = 15.92$, $P < 0.01$, Friedman test; $P$s $< 0.0335$ for stimulation vs. robot and post-stimulation vs. robot sessions comparisons, Dunn’s test), but not the pre-stimulation session. *, **, and *** denote $P < 0.05$, $P < 0.01$, and $P < 0.001$ compared to the robot session, respectively. (G) Stim pairs had higher correlated firing than non-stim pairs during the 20-40 ms post-fire period ($U = 32$, $P = 0.011$; Mann-Whitney U test) but lower during the 60-100 ms window ($U$s $< 37$, $P$s $< 0.023$; Mann-Whitney U test). *

denotes $P < 0.05$ compared to the non-stim pairs. (H) CC peaks of stim pairs during the robot session tended to be higher than those of non-stim pairs ($U = 43.0$, $P = 0.051$; Mann-Whitney U test).

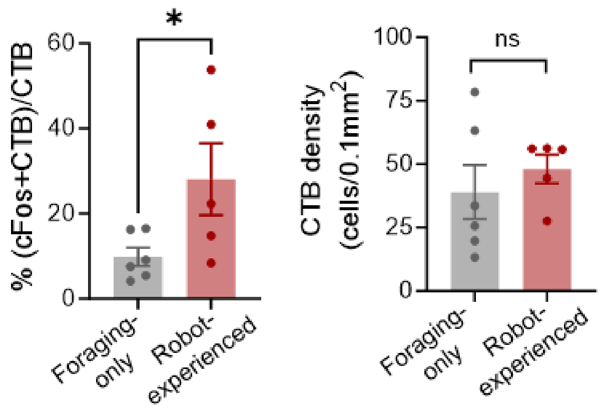


**Figure S5. Double labelling of CTB and c-FOS in PVT cells.** Robot exposure increased the percentage of CTB-labelled PVT neurons expressing c-Fos ($t(9) = 2.278$, $P = 0.0487$). CTB density levels remained similar between groups ($t(9) = 0.7121$, $P = 0.4945$).

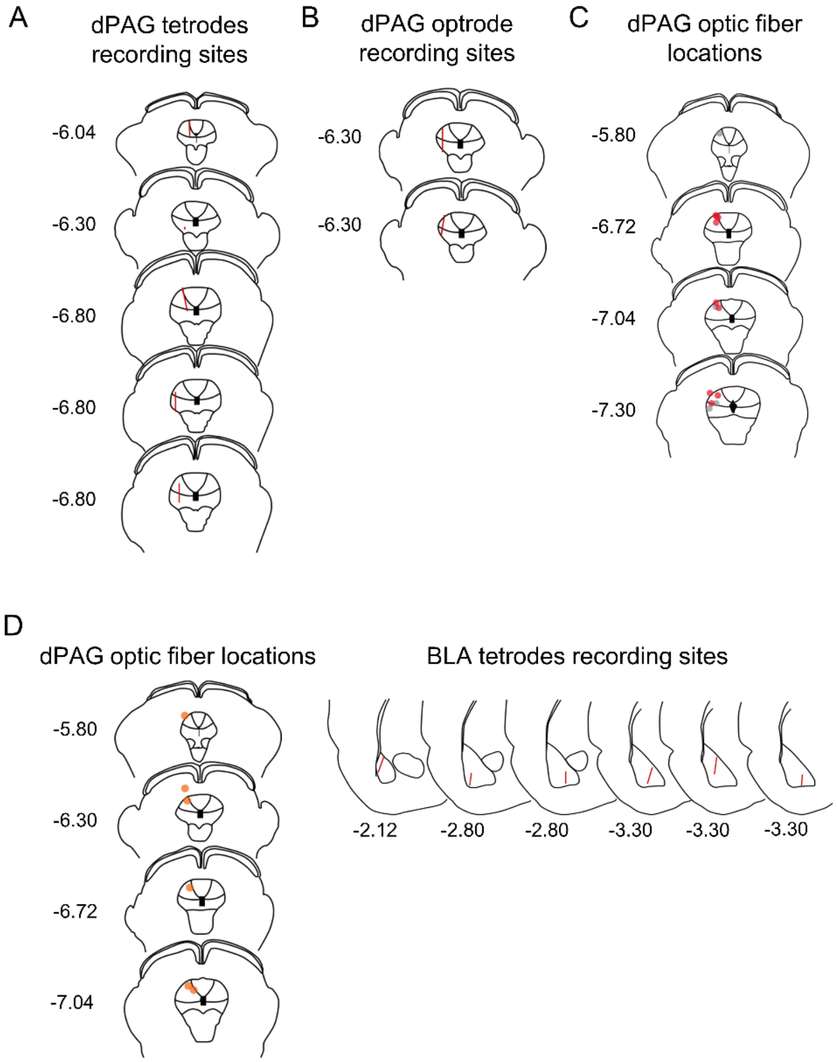


**Figure S6. Histological reconstructions of recording sites in the dPAG and BLA, and optic fiber locations in the dPAG.** (A) Red bars show the trajectory of tetrode recording sites in the dPAG. (B) Red bars depict the trajectory of optrode recording sites in the dPAG. (B) Red and gray circles represent the optic fiber locations for ChR2 and EYFP rats, respectively. (D) Orange circles and red bars indicate the optic fiber locations in the dPAG (*left*) and recording trajectories in the BLA (*right*), respectively. Numerical values represent AP coordinates relative to Bregma.

### Table S1. Normality test results

| Fig | Variables | KS distance | P | Passed (α = 0.01)? | Fig | Variables | KS distance | P | Passed (α = 0.01)? |
|---|---|---|---|---|---|---|---|---|---|
| 1C | Outbound foraging time (s) | | | | 4H | CTB density (% of control) | | | |
| | pre | 0.2916 | <0.0001 | No | | Foraging only | 0.2191 | >0.1000 | Yes |
| | robot | 0.1713 | 0.0033 | No | | Robot experienced | 0.3914 | 0.0118 | Yes |
| | post | 0.2807 | <0.0001 | No | | (cFos+CTB)/CTB (% of control) | | | |
| 1D | Success rate (%) | | | | | Foraging only | 0.267 | >0.1000 | Yes |
| | pre | 1 | <0.0001 | No | | Robot experienced | 0.1972 | >0.1000 | Yes |
| | robot | 0.5102 | <0.0001 | No | S1D | Average firing (spikes/s) | | | |
| | post | 1 | <0.0001 | No | | Pre-robot | 0.3185 | <0.0001 | No |
| 1F | PAG robot cells firing (Z) | | | | | Robot | 0.3045 | <0.0001 | No |
| | pre_bin1 (0-1 s) | 0.1094 | >0.1000 | No | | Post-robot | 0.3197 | <0.0001 | No |
| | pre_bin2 (1-2 s) | 0.2252 | 0.005 | No | | Max firing (spikes/s) | | | |
| | pre_bin3 (2-3 s) | 0.2933 | <0.0001 | No | | Pre-robot | 0.2578 | 0.0005 | No |
| | pre_bin4 (3-4 s) | 0.2553 | 0.0006 | No | | Robot | 0.1969 | 0.0263 | Yes |
| | pre_bin5 (4-5 s) | 0.2324 | 0.0031 | No | | Post-robot | 0.2103 | 0.0125 | Yes |
| | robot_bin1 (0-1 s) | 0.2035 | 0.0184 | Yes | S3H | AUC (Z, 0-0.5s) | | | |
| | robot_bin2 (1-2 s) | 0.2038 | 0.0181 | Yes | | Stim cells | 0.2519 | <0.0001 | No |
| | robot_bin3 (2-3 s) | 0.1826 | 0.0544 | Yes | | nonStim cells | 0.1462 | >0.1000 | Yes |
| | robot_bin4 (3-4 s) | 0.1913 | 0.0353 | Yes | S3I | AUC (Z, 0-5s) | | | |
| | robot_bin5 (4-5 s) | 0.1564 | >0.1000 | Yes | | Stim cells | 0.2161 | <0.0001 | No |
| | post_bin1 (0-1 s) | 0.1948 | 0.0294 | Yes | | nonStim cells | 0.1709 | 0.0799 | Yes |
| | post_bin2 (1-2 s) | 0.153 | >0.1000 | Yes | S4B | CC AUC (Z) | | | |
| | post_bin3 (2-3 s) | 0.0917 | >0.1000 | Yes | | pre | 0.2343 | 0.0008 | No |
| | post_bin4 (3-4 s) | 0.1499 | >0.1000 | Yes | | stim | 0.2587 | <0.0001 | No |
| | post_bin5 (4-5 s) | 0.1822 | 0.0556 | Yes | | post | 0.2898 | <0.0001 | No |
| 2G-J | Control group n (=4) was too small to analyze normality. | | | | | robot | 0.1273 | >0.1000 | Yes |
| | Nonparametric statistics were used for analyzing the data. | | | | S4C | CC peak (Z) | | | |
| 3C | Outbound foraging time (s) | | | | | pre | 0.1998 | 0.009 | No |
| | pre | 0.1481 | 0.0002 | No | | stim | 0.2359 | 0.0007 | No |
| | robot | 0.1155 | 0.0119 | Yes | | post | 0.3407 | <0.0001 | No |
| | post | 0.1483 | 0.0002 | No | | robot | 0.1926 | 0.0142 | Yes |
| 3D | Success rate (%) | | | | S4E | CC AUC (Z) | | | |
| | pre | 1 | <0.0001 | No | | pre | 0.2408 | 0.0138 | Yes |
| | robot | 1 | <0.0001 | No | | stim | 0.2842 | 0.0012 | No |
| | post | 1 | <0.0001 | No | | post | 0.1278 | >0.1000 | Yes |
| 3G | (Outbound foraging time (s) | | | | | robot | 0.2174 | 0.0417 | Yes |
| | pre | 0.1375 | >0.1000 | Yes | | CC peak (Z) | | | |
| | stim | 0.1758 | >0.1000 | Yes | | pre | 0.2297 | 0.0237 | Yes |
| | post | 0.2018 | 0.0808 | Yes | | stim | 0.2183 | 0.0401 | Yes |
| | robot | 0.2389 | 0.0152 | Yes | | post | 0.3022 | 0.0004 | No |
| 3M | Relative firings (Z) - All significant pairs | | | | | robot | 0.2014 | 0.0821 | Yes |
| | pre | 0.1998 | 0.009 | No | S4F | CC AUC (Z) | | | |
| | stim | 0.2359 | 0.0007 | No | | pre | 0.2647 | 0.0454 | Yes |
| | post | 0.3407 | <0.0001 | No | | stim | 0.2759 | 0.0298 | Yes |
| | robot | 0.1926 | 0.0142 | Yes | | post | 0.3317 | 0.0026 | No |
| 3N | Relative firings (Z) - Stim pairs | | | | | robot | 0.1438 | >0.1000 | Yes |
| | pre | 0.2297 | 0.0237 | Yes | | CC peak (Z) | | | |
| | stim | 0.2183 | 0.0401 | Yes | | pre | 0.2153 | >0.1000 | Yes |
| | post | 0.3022 | 0.0004 | No | | stim | 0.2321 | >0.1000 | Yes |
| | robot | 0.2014 | 0.0821 | Yes | | post | 0.4233 | <0.0001 | No |
| | Relative firings (Z) - nonStim pairs | | | | | robot | 0.3327 | 0.0025 | No |
| | pre | 0.2153 | >0.1000 | Yes | S4G | CC AUC (Z) | | | |
| | stim | 0.2321 | >0.1000 | Yes | | stim pair (0-20 ms) | 0.1021 | >0.1000 | Yes |
| | post | 0.4233 | <0.0001 | No | | stim pair (20-40 ms) | 0.2242 | 0.0307 | Yes |
| | robot | 0.3327 | 0.0025 | No | | stim pair (40-60 ms | 0.192 | >0.1000 | Yes |
| 4C | Latency to procure pellets (s) | | | | | stim pair (60-80 ms) | 0.2829 | 0.0013 | No |
| | Base_Foraging only | 0.3539 | 0.0181 | Yes | | stim pair (80-100 ms) | 0.2552 | 0.0065 | No |
| | Base_Robot experienced | 0.2801 | 0.0253 | Yes | | nonstim pair (0-20 ms) | 0.1589 | >0.1000 | Yes |
| | Test_Foraging only | 0.2047 | >0.1000 | Yes | | nonstim pair (20-40 ms) | 0.2269 | >0.1000 | Yes |
| | Test_Robot experienced | 1 | <0.0001 | No | | nonstim pair (40-60 ms) | 0.2308 | >0.1000 | Yes |
| 4E | Fos-positive cells (% of control) | | | | | nonstim pair (60-80 ms) | 0.3189 | 0.0048 | No |
| | PVT_Foraging only | 0.2505 | >0.1000 | Yes | | nonstim pair (80-100 ms) | 0.2742 | 0.0319 | Yes |
| | PVT_Robot experienced | 0.3508 | 0.0010 | No | S4H | CC peak | | | |
| | IMD_Foraging only | 0.2808 | >0.1000 | Yes | | Stim pairs | 0.3327 | 0.0025 | No |
| | IMD_Robot experienced | 0.1831 | >0.1000 | Yes | | nonStim pairs | 0.2014 | 0.0821 | Yes |
| | CM_Foraging only | 0.2105 | >0.1000 | Yes | S5 | CTB density (cells/0.1 $mm^2$) | | | |
| | CM_Robot experienced | 0.2463 | 0.0866 | Yes | | Foraging only | 0.2488 | >0.1000 | Yes |
| | Rh_Foraging only | 0.2892 | >0.1000 | Yes | | Robot experienced | 0.3321 | 0.0748 | Yes |
| | Rh_Robot experienced | 0.1480 | >0.1000 | Yes | | % (cFos+CTB)/CTB | | | |
| | Re_Foraging only | 0.1952 | >0.1000 | Yes | | Foraging only | 0.2206 | >0.1000 | Yes |
| | Re_Robot experienced | 0.1330 | >0.1000 | Yes | | Robot experienced | 0.2192 | >0.1000 | Yes |
